# A NanoBRET-based assay monitoring interactions in the USP18 signaling hub identifies the first cell-penetrant small molecule compromising USP18/ISG15 binding

**DOI:** 10.1101/2025.03.30.646166

**Authors:** Sandra Hess, Marta Campos Alonso, Michael Brand, Susan Lauw, Urs Lindenmann, Katharina Göbel, Paul P. Geurink, Günter Fritz, Rainer Riedl, Klaus-Peter Knobeloch

## Abstract

Protein modification by interferon-stimulated gene 15 (ISG15), termed ISGylation, exhibits antiviral properties and influences tumorigenesis, genome stability and metabolic processes. ISGylation is counteracted by the specific protease USP18. Likewise, viral proteases such as the papain-like protease (PLpro) from SARS-CoV-2 cleave ISG15 to undermine the host immune response. Beyond its role as a deISGylating enzyme, USP18 acts as a major negative regulator of the IFN signaling pathway in a STAT2-dependent manner. In humans, unconjugated ISG15 secures USP18 stability and the absence of USP18 or impaired STAT2/USP18 binding cause fatal interferonopathies. Thus, the USP18 signaling hub represents a critical checkpoint for type I IFN signaling and ISGylation, qualifying it as a promising immune and cancer drug target. However, suitable assays to monitor protein-protein interactions (PPIs) within the USP18/ISG15/STAT2 signaling hub and to screen for PPI modulators are missing and no specific inhibitors targeting USP18 interactions are available.

To address this gap, we developed a method based on the NanoLuc luciferase (NLuc) Bioluminescence Energy Transfer (NanoBRET) assay system to study PPIs. Firstly, we generated stable cell lines suitable to monitor USP18/ISG15 and USP18/STAT2 interactions, providing a semi high-throughput screening (HTS)-compatible platform. In combination with a virtual pre-screen of 60,000 compounds against USP18 *in silico*, this assay allowed us to identify a first small molecule (ZHAWOC8655) that compromises cellular USP18/ISG15 binding and inhibits USP18 protease activity *in vitro*. To further explore the potential of using the NanoBRET system for testing PPI modulators, we evaluated the effect of GRL0617, a compound which was shown to disrupt the interaction between SARS-CoV-2 PLpro/ISG15 as well as SARS-CoV-2 PLpro/ubiquitin. NanoBRET based stable cell lines as presented here will be suitable for monitoring PPIs in other multiprotein complexes after various stimuli, mutations or small molecule administration and can be challenged with siRNA or CRISPR/Cas9 libraries to identify previously unrecognized regulators.

## Introduction

Type I Interferons (IFNs) are essential cytokines in the innate immune response and, as such, critical for the defense against pathogens and neoplasia(1,2). IFNs bind to the ubiquitously expressed type I IFN receptor (IFNAR), which is composed of two transmembrane subunits, IFNAR1 and IFNAR2, and signals via the JAK/STAT pathway that results in the transcription of hundreds of ISGs with far-reaching immunomodulatory effects and direct antimicrobial and anti-neoplastic effects. By a complex regulatory network, type I IFNs link the innate with the adaptive immune response(3,4). Therefore, consistent with its central role, the IFN response requires complex and balanced regulation to prevent pathological situations originating from immune dysfunction, autoimmunity or hyperinflammation.

Consequently, IFNs, IFN signaling and IFN effector systems represent attractive targets for therapeutic intervention strategies. IFN-α and IFN-β are currently used for the treatment of chronic hepatitis B, hairy cell leukemia, chronic myeloid leukemia and malignant melanoma(5). Conversely, overshooting IFN responses are addressed by anti-IFNAR antibodies and JAK inhibitors(6). However, there are fundamental limitations in current therapeutic approaches such as side effects, desensitization and relapses(6). Thus, there is a demand for novel modulators of the IFN response targeting previously unaddressed proteins, which could also be combined with existing drugs to achieve synergistic effects or overcome resistance.

ISG15 represents one of the genes most strongly upregulated upon type I IFN signaling(7). In a cytokine-like manner, free ISG15 can be secreted from cells and bind to the lymphocyte function-associated antigen-1 (LFA-1), promoting IFN-γ secretion from natural killer (NK) and T cells(8). Moreover, ISG15 acts as an ubiquitin-like modifier and is covalently linked to target proteins, in a process called ISGylation. Structurally, ISG15 is composed of two ubiquitin-like domains. In analogy to ubiquitin modification, ISGylation is mediated by the consecutive action of an E1 activating enzyme (Ube1L), E2 conjugating enzyme (UbcH8) and several E3 ligases like *m*HERC6, *h*HERC5, HHARI and EFP. ISGylation was shown to occur in a co-translational manner, favoring modification of highly expressed viral proteins in infected cells, which in turn interferes with virus assembly or function(9,10). In addition, cellular proteins involved in antiviral defense or the export of viral particles were also shown to be ISGylated, contributing to the antiviral function of ISG15(11).

ISGylation is reversed by ubiquitin-specific protease 18 (USP18). Structural and functional investigations on USP18 alone and in complex with ISG15 by X-ray crystallography showed that USP18 is a specific ISG15 isopeptidase (12,13). Selective inactivation of only the protease function of USP18 in mice (USP18^C61A/C61A^) stabilized ISGylation and resulted in increased resistance against viral infections(14,15). Pathogens have evolved proteases capable of deconjugating ISG15. An example is the PLpro enzyme from SARS-CoV-2, which exhibits deISGylating activity, thereby contributing to the strong immune evasion properties of the virus(16). Morever, ISGylation suppresses breast cancer growth and metastasis by affecting the tumor microenvironment and inactivation of the deISGylating activity of USP18 reduced tumor burden in a mouse breast cancer model(17).

Remarkably, beside its function as the main ISG15 protease, USP18 represents one of the major negative regulators of the type I IFN signaling pathway(18–20). On the one hand, USP18 has been shown to interact with the IFNAR subunit 2, competing with JAK1 for receptor binding and thereby counteracting proximal signaling(21). On the other hand, USP18 has been reported to interfere with IFN-induced dimerization of subunits 1 and 2 of IFNARupon IFN binding. It was shown that STAT2 is an essential adapter in USP18-mediated suppression of type I IFN signaling by recruiting USP18 to IFNAR2 via its constitutive membrane-distal STAT2-binding site(22,23). These data show that USP18 is a bifunctional protein and exerts this important function irrespective of the ISG15 protease activity via interaction with aforementioned binding partners.

USP18’s non-enzymatic function as a negative regulator of IFN signaling is essential to prevent uncontrolled microglia activation and lack of the protein causes interferonopathy in mice (19). USP18 also plays a major role in IFN desensitization where an initial injection of IFN causes unresponsiveness of the IFN signaling pathway after re-challenge (24). Interestingly, USP18-deficient mice do not exhibit IFN-desensitization uncovering USP18 as a potential target to restore the IFN response in clinical regiments(18). Concordantly, silencing of USP18 in human cells potentiates the antiviral activity of IFN against hepatitis C virus infection(25). Free ISG15 stabilizes USP18 in humans by counteracting proteasomal degradation thereby ensuring its negative regulatory capacity. Accordingly, human patients lacking ISG15 exhibited decreased levels of USP18, increased IFN signaling and interferonopathy(26). The clinical relevance of USP18 function is further highlighted by the observation that genomic loss of functional USP18 in humans causes embryonic lethality and perinatal death associated with aberrant IFN signaling and brain calcification(20) or inflammation(27). Likewise, mutations in STAT2 causes human interferonopathy in a USP18-dependent manner(22,28).

The central role of USP18 as a deISGylase and as a main checkpoint for IFN signaling has raised interest in therapeutically targeting its protein interactions and/or catalytic activity (29). The unavailability of sufficient amounts of recombinant enzymatically active human USP18 has hampered classical *in vitro* high-throughput screening approaches. As IFN regulatory properties of USP18 are mediated via interaction with protein partners, small molecule modifiers for these PPIs cannot be identified by classical enzymatic assay systems. Thus, there is an unmet need for a screening compatible assay that can detect alterations in USP18 protein-protein interactions (PPIs). Likewise, investigations on USP18 PPIs, competition among different binding partners, the consequences of particular mutations and the response to extracellular stimuli would benefit from such a tool.

Bioluminescence Resonance Energy Transfer (BRET) assays have been successfully used to study PPIs in a semi-quantitative manner in intact cells. In this system, NanoLuc luciferase (NLuc) is fused to protein A, whereas the binding partner B is linked to a modified haloalkane dehalogenase moiety (HaloTag) designed to covalently bind to synthetic ligands (HaloTag ligands). When proteins A and B interact, an added fluorescently labelled HaloTag ligand is excited by the NLuc. This requires close proximity (less than 10 nm) between the interacting proteins and allows calculation of a BRET ratio (the ratio of NLuc signal to the fluorescent signal) as a readout of PPI. The BRET ratio allows to quantitatively monitor alterations in PPI, e.g. upon administration of interfering small molecules, application of exogenous stimuli, insertion of mutations in protein A or B or expression of unlabeled binding partners(30). However, the use of the BRET system as a PPI assay requires considerable adaptation for individual interacting partners and has rarely been reported for applications in screening for small molecule inhibitors.

Here we developed such a BRET-based assay for USP18 and its interaction partners ISG15 and STAT2. By generating stable cell lines with the BRET components, the assays were optimized to be suitable for semi high-throughput screening (HTS) in a 384-well format. To further validate that such a systems can be applied to detect small molecules interfering with ISG15-protease function, we established a BRET assay for the deISGylase SARS-CoV-2 PLpro and ISG15 and validated that PPI is compromised by an established SARS-CoV-2 PLpro inhibitor. Subsequently, the stable USP18/ISG15 BRET cell line was used to screen a targeted compound library preselected against the USP18/ISG15 interaction by *in silico* screening with AutoDock Vina(31). Using this approach, we identified, compounds capable of inhibiting the ISG15/USP18 interaction in a cellular context and USP18 catalytic activity *in vitro*.

## Results

### Development of a NanoBRET-based assay to monitor PPIs of USP18 with ISG15

The bioluminescence resonance energy transfer assay NanoBRET developed by Promega excels other BRET assays due to its enhanced signal stability, increased sensitivity and extended dynamic range, enabling more precise and reliable PPI monitoring in live-cell contexts. Although this assay has been used in several studies, it has typically been employed in transient settings requiring plasmid transfections, which can lead to variable expression levels, limited reproducibility and the inability to conduct long-term experiments. To overcome these limitations, we decided to develop stable cell lines using the Promega NanoBRET assay to monitor PPIs within the USP18/ISG15/STAT2 signaling hub.

As a prerequisite for stable cell line development, we first established and optimized the standard NanoBRET assays. Therefore, the coding sequences of human USP18 (hUSP18) and human ISG15 (hISG15) were cloned into expression vectors to generate fusion constructs with the donor NLuc and the acceptor HaloTag. Importantly, the geometric orientation of the donor and acceptor relative to the interaction interface of the target proteins significantly influences the efficiency of energy transfer. Additionally, the Nluc and the relatively large HaloTag may interfere with protein interactions due to steric hindrance. For ISG15, only N-terminal fusions were feasible due to its precursor protein undergoing C-terminal processing(32). Therefore, various configurations of the donor and acceptor fusions were tested. As depicted in the workflow, the fusion vectors were co-transfected and the NanoBRET assay was subsequently performed (Figure 1B).

**Figure 1:**
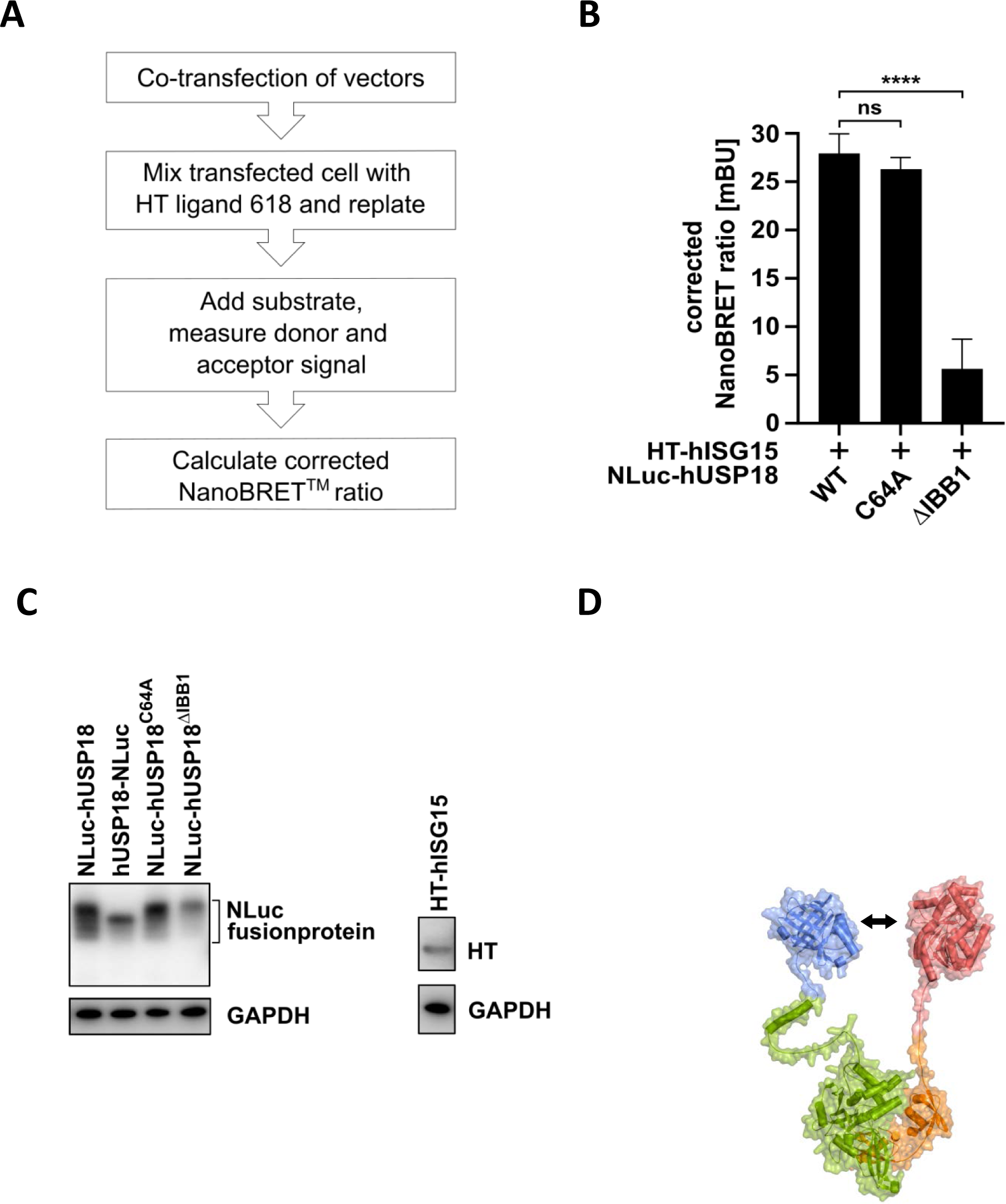
Establishing a PPI assay for hUSP18 and hISG15. **(A)** Schematic workflow of the NanoBRET™ assay (right side). **(B)** Comparison of the NanoBRET™ ratio of the interaction of hISG15 with hUSP18WT, USP18C64A or hUSP18ΔIBB1. Data from at least six independent experiments. **(C)** Immunoblot analysis of the depicted proteins using NLuc®, HaloTag® (HT) and GAPDH specific antibodies. **(D)** Schematic representation of the PPI assay (NanoBRET™) for hUSP18 and hISG15. Donor NLuc® luciferase was N-terminally coupled to USP18 and ISG15 to the acceptor HaloTag® (left side). The oxidation of the Luciferase substrate furimazine enables energy transfer to the acceptor molecule attached to the HaloTag if the two proteins are in close proximity (less than 10 nm). Respective structure models provide evidence that Nluc- or HaloTag-fusion do not sterically hinder USP18-ISG15 interaction. All data presented in mean ± standard deviation. Statistical analysis was performed using one-way ANOVA followed by Bonferroni’s multiple comparison test (****P < 0.0001; ns, not significant).

A robust interaction between NLuc fused to the N-terminus of USP18 and HaloTag fused to the N-terminus ISG15 was observed, indicated by the NanoBRET ratio. Notably, this interaction was preserved with the catalytically inactive USP18 mutant (hUSP18^C64A^), as expected. As a negative control for this PPI pair, we introduced mutations analogous to those identified in previous structural studies of the mUSP18/mISG15 complex(13). These studies highlighted a hydrophobic region of USP18, termed ISG15-binding box 1 (IBB1), as critical for the interaction. Accordingly, we replaced Ala141, Leu145, His255 and Pro195 in hUSP18 with the corresponding residues from USP17 to generate the hUSP18^ΔIBB1^ variant, aiming to disrupt the interaction between hUSP18 and hISG15. As expected, the hUSP18^ΔIBB1^ construct displayed a markedly reduced NanoBRET ratio compared to hUSP18WT (Figure 1C), validating the effectiveness of the assay in detecting and modulating hUSP18/hISG15 interactions. To reduce the risk of false positive interaction signals due to non-specific energy transfer from donor to acceptor, the ratio of NLuc-hUSP18 to HaloTag-hISG15 vector was optimized to 1:1000 (data not shown) as suggested by the manufacturer’s protocol. Expression of the selected vector constructs was verified by Immunoblotting using antibodies specific against either NLuc (USP18 and its variants) or HaloTag (ISG15) (**Figure 1D**). Moreover, structural modelling of the fusion proteins suggested that neither NLuc nor HaloTag introduced steric hindrance to the interaction (**Figure 1A**).

### Establishment of a NanoBRET assay to monitor hUSP18/hSTAT2 protein-protein interaction

It has been shown that the interaction of hUSP18 with hSTAT2 is essential to secure the negative regulatory function of hUSP18 on type I IFN signaling(23). Concordantly, patients carrying mutations in hSTAT2 which compromise hUSP18/hSTAT2 interactions have been reported to suffer from type I interferonopathies(22,28). We therefore also established BRET-based assays for monitoring the interaction between hUSP18 and hSTAT2. Structural models of the fusion proteins put forward that neither the NLuc nor the HaloTag cause steric hindrance of USP18/STAT2 interactions when the full-length hSTAT2 coding sequence was C-terminally fused with the HaloTag and combined with N-terminal NLuc-hUSP18 (Fi**gure 2A**). NanoBRET assay was performed as described above. Like for hISG15, replacement of hUSP18^WT^ with the catalytically inactive variant (hUSP18^C64A^) did not affect the PPI to hSTAT2. Expression of the hSTAT2-HaloTag fusion protein was detected by an anti-HaloTag antibody (**Figure 2C**).

**Figure 2:**
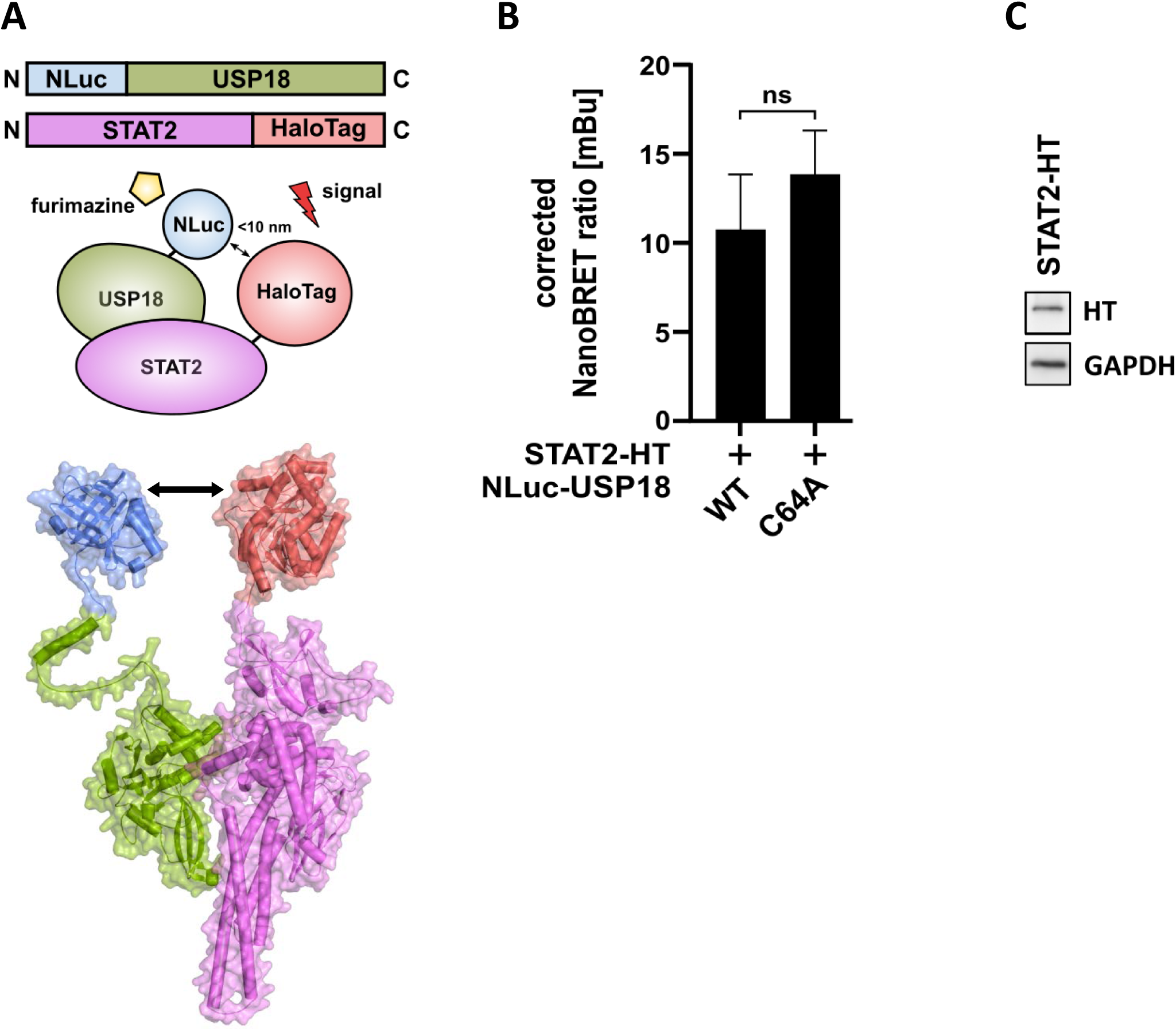
Establishing a protein-protein interaction assay for human USP18 and STAT2. **(A)** Schematic depiction of the NanoBRET assay constructs and respective structural models for USP18 and STAT2. The NanoLuc (NLuc) Luciferase was N-terminally coupled to USP18 and HaloTag C-terminally to STAT2. **(B)** NanoBRET assay for the interaction of USP18^WT^/STAT2 and USP18^C64A^/STAT2. Data from at least 3 independent experiments. Depicted in mean ± stdev. Statistical analysis was performed using one-way ANOVA followed by Bonferronis multiple comparison test (\*\*\*\**P* < 0.0001), ns=not significant. **(C)** Immunoblot analysis of the STAT2-HT fusionprotein. HEK293T cells were transfected with 2 µg DNA of the indicated construct. STAT2-HaloTag was detected with anti-HaloTag antibody. GAPDH was used as loading control.

### Adapting the developed BRET-based assay into a screening platform to identify compounds modulating PPIs

Once NanoBRET assays monitoring PPIs within the USP18/ISG15/STAT2 signaling hub were established, we wondered whether these systems could be used to identify molecules disrupting these interactions. We decided to first test this hypothesis using the SARS-CoV-2 encoded papain-like protease (SARS-CoV-2-PLpro) which not only removes ISG15 from cellular and viral substrates to undermine the host immune response but also ubiquitin, albeit less efficiently (16). The availability of inhibitory compounds against this protease, along with its similarity to the structure and catalytic function of USP18, made SARS-CoV-2-PLpro an ideal case study to test our proposal. Therefore, we established a NanoBRET-based assay for monitoring the interaction between SARS-CoV-2-PLpro and hISG15 as well as SARS-CoV-2-PLpro and ubiquitin, using a similar strategy as described above. Different fusion variants were tested as depicted in **Figure 3A**. Robust interaction was detected when the SARS-CoV-2-PLPro was fused with the NLuc and expressed together with hISG15 and ubiquitin fused with HaloTag (**Figure 3A**). We next evaluated whether GRL-0617, an established inhibitor of the SARS-CoV-2-PLpro activity, not only blocks the catalytic function of this enzyme but also compromises its interaction with hISG15 and/or ubiquitin. As depicted in **Figure 3B**, increasing amounts of inhibitor diminished the NanoBRET signal of SARS-CoV-2-PLpro/hISG15 and also SARS-CoV-2-PLpro/ubiquitin, indicating that the compound indeed interferes with the interaction of the proteins. In contrast, interactions between MDM2 and p53 as well as USP18 and hISG15 were not affected.

**Figure 3:**
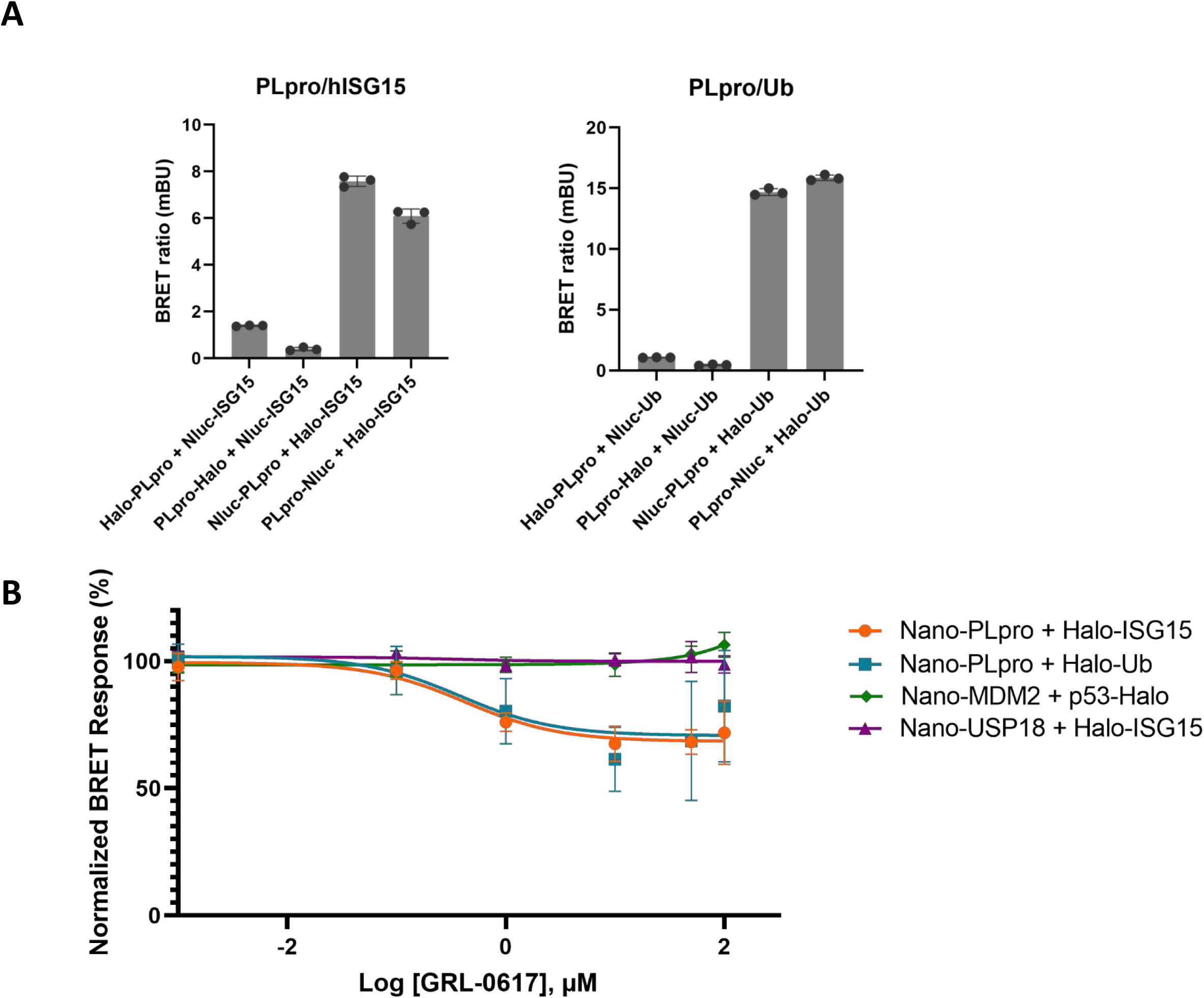
Establishing a protein-protein interaction assay for SARS-CoV-2-PLpro and human ISG15 or ubiquitin. **(A)** Schematic depiction of the optimization of the NanoBRET assay for SARS-CoV-2-PLpro and human ISG15/ubiquitin using different fusion variants. **(B)** NanoBRET assay for SARS-CoV-2-PLpro and human ISG15 or ubiquitin with increasing amounts of the SARS-CoV-2-PLpro inhibitor GRL-0617 and respective controls (MDM2/p53 and USP18/ISG15) as indicated. NanoBRET signals were normalised to DMSO control.

These results not only validate the specificity and sensitivity of the NanoBRET assay to monitor alterations in PPIs, but also provide a proof of principle that respective NanoBRET assays are applicable for the identification or validation of compounds interfering with substrate recognition of deISGylases or deubiquitinating enzymes.

### Adaptation of NLuc-hUSP18/HaloTag-hISG15 and NLuc-hUSP18/hSTAT2-HaloTag NanoBRET systems to semi HTS compatible formats

As previously mentioned, the NanoBRET assay delivers exceptional signal stability, sensitivity and dynamic range, making it ideal for precise PPI monitoring in cells. However, transient expression systems are hindered by inconsistent expression levels and limited reproducibility, which poses challenges, especially when optimizing the assay for semi HTS applications with restricted cell quantities per vial. Furthermore, for each testing round cells would need to be transfected prior to addition of the compound library, which adds additional variations and promotes inconsistencies. To circumvent these problems, we aimed to generate cell lines where the aforementioned NanoBRET components are stably integrated into the genome.

As the amount of NLuc expression needs to be limited (which is achieved in the transient systems by transfecting reduced amounts of the NLuc-hUSP18 vector), we made use of a vector with a bidirectional promotor as schematically depicted in **Figure 4A**. For details see Suppl Figures 1-3 and material and methods section. Here, expression of NLuc-hUSP18 is driven by a minimal CMV promoter combined with a full-length CMV promoter driving expression of either HaloTag-hISG15 (**Figure 4B**) or hSTAT2-HaloTag fusion proteins (**Figure 4C**). Moreover, to generate a control cell line, NLuc-hUSP18^ΔIBB1^ was used along with HaloTag-hISG15. As schematically depicted in **Figure 4A**, the final vectors were transfected and stably integrated into the genome of HEK293 cells. Bulk cell PPI analysis by NanoBRET assay revealed that this approach is well suited to monitor hISG15/hUSP18 (**Figure 4B**) and hUSP18/hSTAT2 interactions (**Figure 4C**). As expected, no NanoBRET signal is detected in the hISG15/hUSP18^ΔIBB1^ negative control assay (**Figure 4B**). Although expression levels of the NanoBRET components are mostly determined by the delivered bidirectional promotor, local effects of the integration site such as chromatin accessibility or modification status might differ between individual cells. To further minimize variability and ensure robust NanoBRET signals, individual cell clones were isolated, expanded and tested for the suitability to monitor PPI. As shown in **Figure 4D/E** NanoBRET measurements revealed that these single cell derived populations are functional in the NanoBRET assay to monitor respective PPI pairs. Using this strategy, we managed to establish cell-based NanoBRET assays for hUSP18/hISG15 and hUSP18/hSTAT2, which can robustly be performed in a 384 well format and are thus applicable for semi HTS approaches.

**Figure 4:**
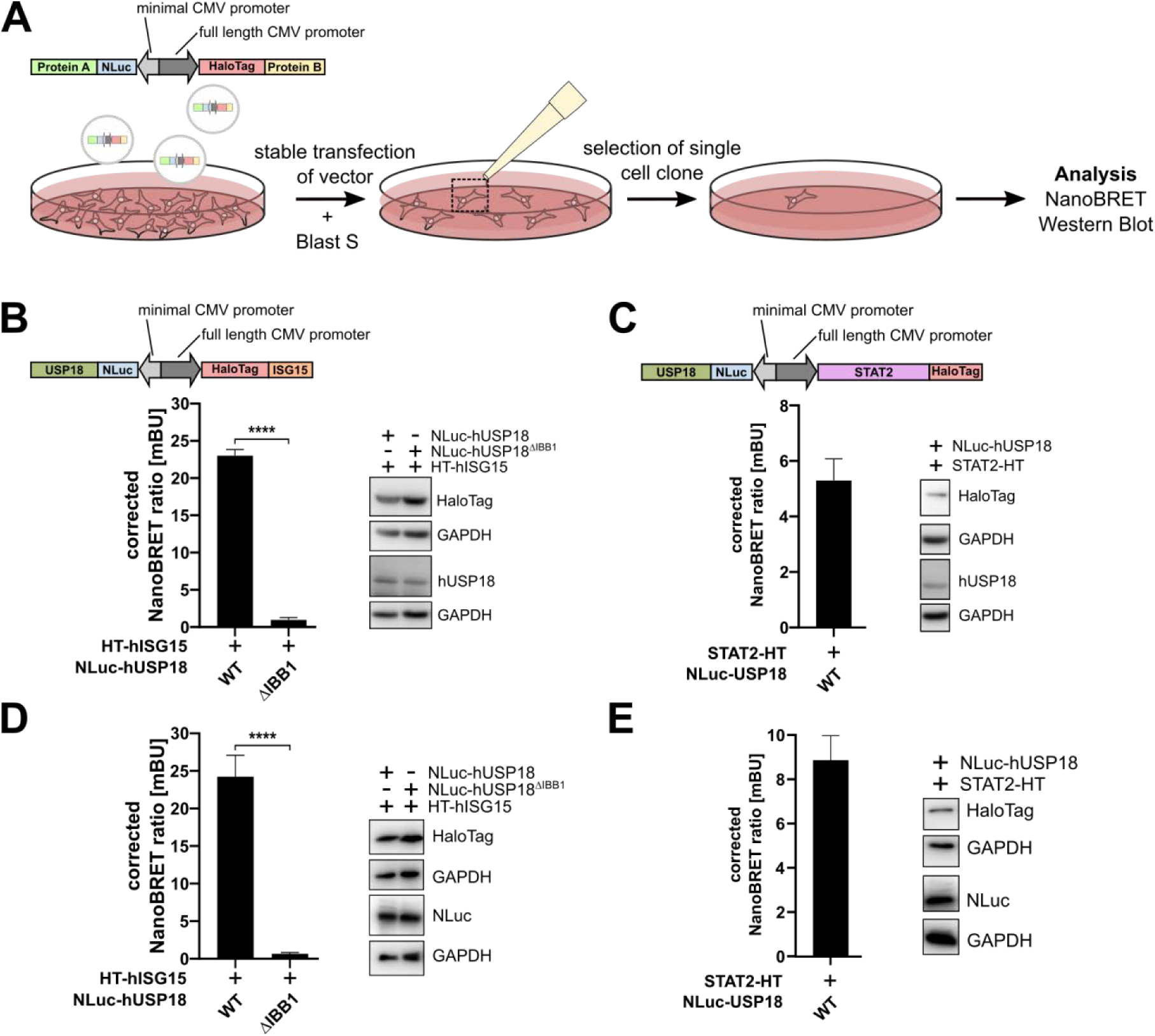
Generation of stable cell lines for the protein-protein interaction of human USP18 with human ISG15 and human STAT2. **(A)** Schematic depiction of the experimental approach used to generate the stable NanoBRET cell lines. A vector with a biased bi-directional CMV promotor system (BiBRET from Promega) was used for the expression, ensuring a higher HaloTag-fusion protein expression as compared to NanoLuc-fusion protein. HEK293 cells were transfected with the linearized vector. Successful integration was screened using antibiotic selection with Blasticidin S (Blast S). Single cell clones were selected and used for all further experiments. **(B/C)** NanoBRET and Immunoblot analysis of the antibiotic resistant cell pool (bulk transfection) of the HEK293 cells transfected with NanoLuc N-terminally coupled to human USP18^WT^ or USP18^ΔIBB1^ and HaloTag to human ISG15 (B) or human STAT2 **(C)**. **(D/E)** NanoBRET and Immunoblot analysis of the antibiotic resistant selected single cell clones for USP18/ISG15 (D) and USP18/STAT2 expression (E). Data from at least 3 independent experiments. Depicted in mean ± stdev. Statistical analysis was performed using unpaired Student’s t-Test followed (\*\*\*\**P* < 0.0001; \*\*\**P* < 0.001).

### Selection of a focused small molecule library to target the interface of USP18 and ISG15 by virtual screening

The crystal structure of mUSP18 (PDB code: 5cht) and mUSP18 in complex with mISG15 (PDB code: 5chv), provided the first structural insight to this pharmaceutical target(13). The interaction between mISG15 and mUSP18 includes an extended, flat hydrophobic protein surface, making it a difficult target for small molecules. Targeting PPIs has been a long standing and challenging problem(33). Nevertheless, in recent years it was demonstrated that inhibiting PPIs is possible by targeting the region of amino acids critically involved in PPI (hot spot amino acids). For example, the discovery of the allosteric inhibitors GNE6640 and GNE6776 against USP7 by Genentech exemplifies that the shallow protein surface of USPs can indeed be blocked(34,35).

As described above, the structural and mutational studies on USP18 revealed the hot spot interactions between USP18 and its substrate ISG15 to reside in the conserved IBB1 region (including residues Ala138, Leu142, Ser192, His251) and the blocking loop comprising residues 256-263 of mUSP18(13). We envisaged a virtual screening targeting the interface of mUSP18 and mISG15 to identify small molecule PPI disruptors for this protein complex. Commercially available PPI disruptor-like compound libraries from Enamine and Life Chemicals containing 60,000 compounds were screened against USP18 using AutoDock Vina(31) and resulted in 600 virtual hits with a binding score of < −8.2 kcal/mol. After visual inspection of the compounds in the hypothesized binding mode to USP18, 365 promising compounds were selected (**Figure 5A**).

**Figure 5:**
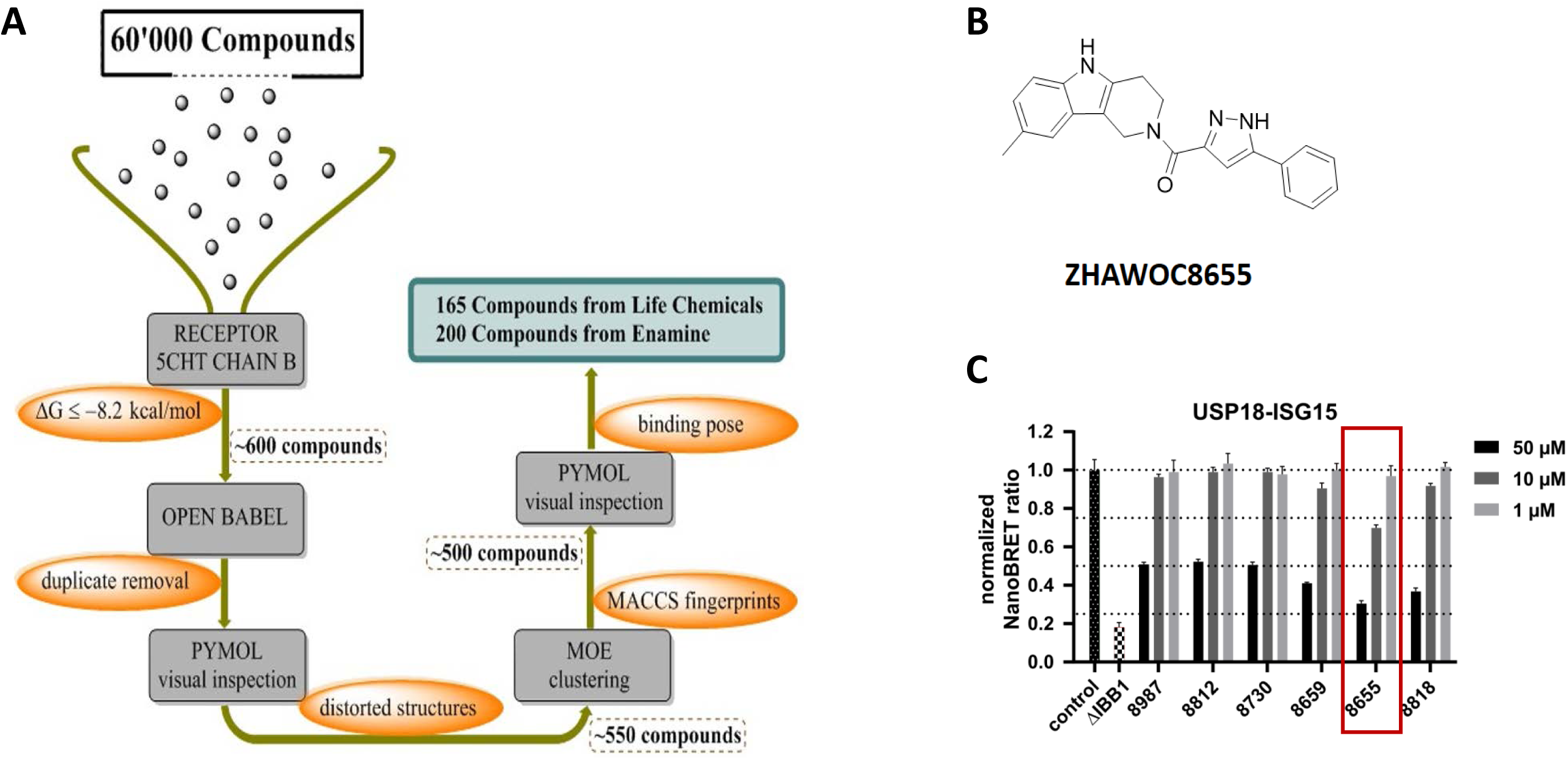
**(A)** Schematic representation of the selection process for the virtual screening of 60’000 compounds from Life Chemicals and Enamine. The compounds were first docked using Autodock Vina (PMID 19499576) to USP18 (PDB: 5cht chain B) and a threshold of −8.2 kcal/mol for ΔG was used to identify 600 compounds which matched this criteria. Duplicates from the remaining compounds were removed using Open Babel (PMID: 21982300). By visual inspection of the binding pose unrealistic distortions and the structures were removed. To remove similar compounds with similar binding pose a clustering using MACCS fingerprints was calculated in Molecular Operating Environment. **(B)** Structure of the identified hit molecule ZHAWOC8655. **(C)** Effect of ZHAWOC8655 (marked in red) and other initial hits on the USP18/ISG15 NanoBRET ratio at indicated concentrations.

### Identification of small molecules interfering with hUSP18/hISG15 interaction

The focused small molecule library was used to identify compounds interfering with USP18/ISG15 PPI using the stable NLuc-hUSP18/HaloTag-hISG15 NanoBRET cell line described above (**Figure 4D**). Each compound was applied at a concentration of 50 µM in the initial screening, which identified a total of six compounds with different scaffolds. Compound ZHAWOC8655 (**Figure 5B**) was selected for further studies as the most promising candidate based on its synthetic accessibility and its dose-dependent inhibition at concentrations of 50, 10 and 1 µM (**Figure 5C**).

To confirm the activity and exclude any possible artefacts caused by impurities, the screening compound ZHAWOC8655 was resynthesized. The synthesized compound was first reevaluated in the BRET assay system against hUSP18/hISG15. In excellent agreement with the previous data, the modulating activity of the resynthesized hit compound ZHAWOC8655 was confirmed and showed a typical dose response curve, validating the results from the initial screen (**Figure 6A**). Of note, the compound did not unspecifically interfere with the NanoBRET assay system, as the p53/MDM2 interaction BRET assay was unaffected by ZHAWOC8655 (**Figure 6B**).

**Figure 6:**
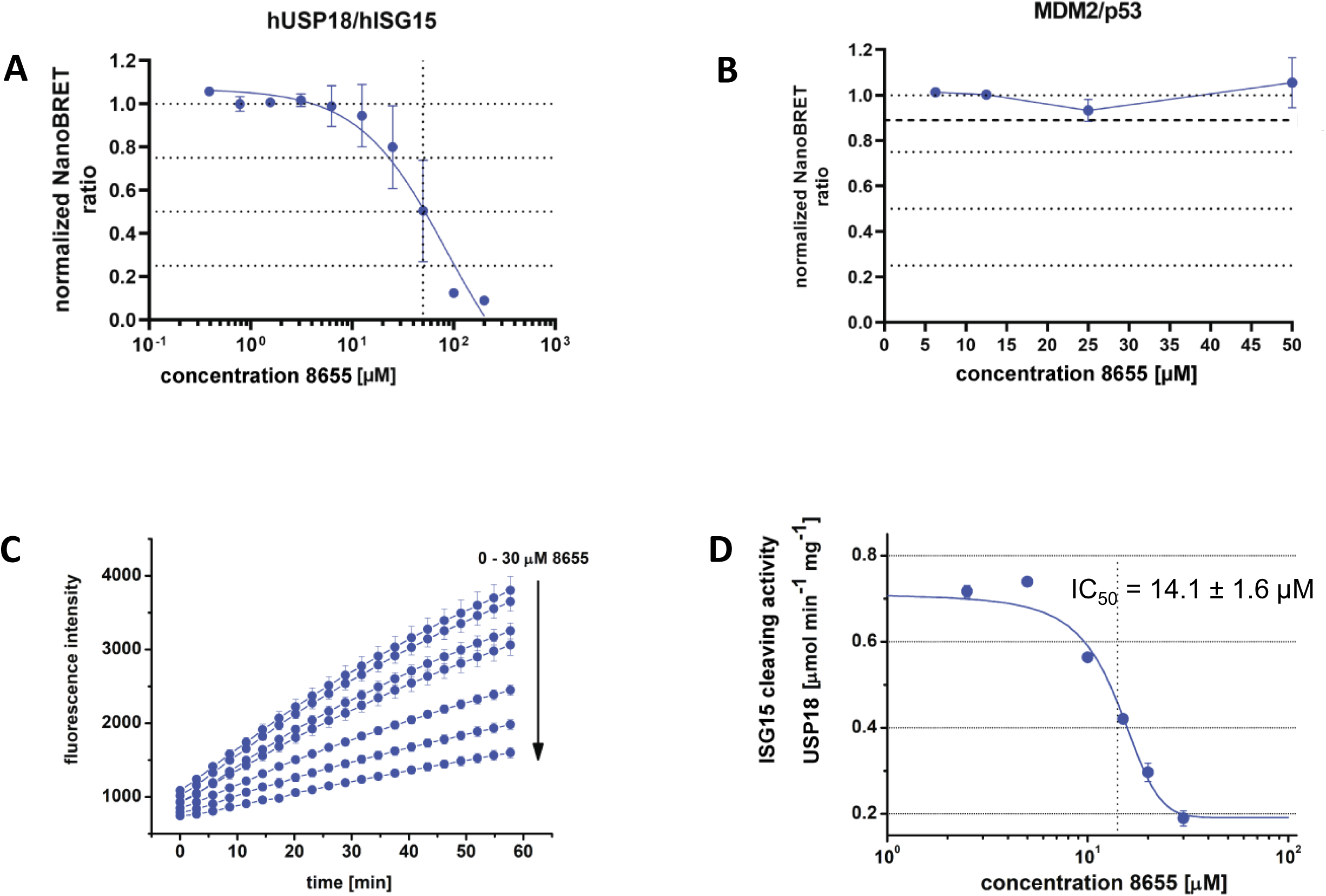
**(A)** The BRET ratio of human USP18/ISG15 interaction decreases in the presence of compound 8655 in a dose dependent manner. **(B)** The interaction of MDM2/p53 is not affected by the compound in the same concentration range **(C)** Inhibition of USP18 catalysed ISG15 cleavage by compound 8655. The left hand panel depicts increase of fluorescence by rhodamine liberated upon cleavage of ISG15-rhodamine. **(D)** Doses respond curve of the specific activity of human USP18 versus the compound concentration.

### The identified USP18/ISG15 PPI-inhibitor ZHAWOC8655 interferes with the deISGylating function of USP18 in vitro

Upon binding to ISG15, USP18 undergoes drastic structural alterations. Most strikingly, the residues forming the catalytic triad essential for proteolytic activity reside in an inactive arrangement in unbound USP18 and are repositioned to form an active triad upon ISG15 binding(13). We thus speculated that the identified inhibitor of hUSP18/hISG15 interaction might affect the protease activity of USP18 that cleaves ISG15 from target proteins. Therefore, we used a well-established model substrate where the C-terminal domain of ISG15 is covalently linked to a derivate of the fluorophore rhodamine. USP18 protease activity can thus be monitored by an increase in fluorescence of the liberated fluorescent dye. As shown in **Figure 6C**, compound ZHAWOC8655 inhibited the enzymatic activity of recombinant human USP18 *in vitro* in a dose dependent manner and analysis of the kinetics revealed an inhibition constant of 14.1 ± 1.6 µM (**Figure 6D**). The docking results depicted in **Figure 7** suggest that compound ZHAWOC8655 binds in close proximity to the IBB-1 site in hUSP18, partially blocking the deep cleft that binds the C-terminus of ISG15 and the hydrophobic plane of the palm region of hUSP18. In the predicted binding pose, the compound forms three hydrogen bonds to the side chains of residues Asp139, Gln142 and Tyr363 as well as hydrophobic interactions with the surrounding residues.

**Figure 7:**
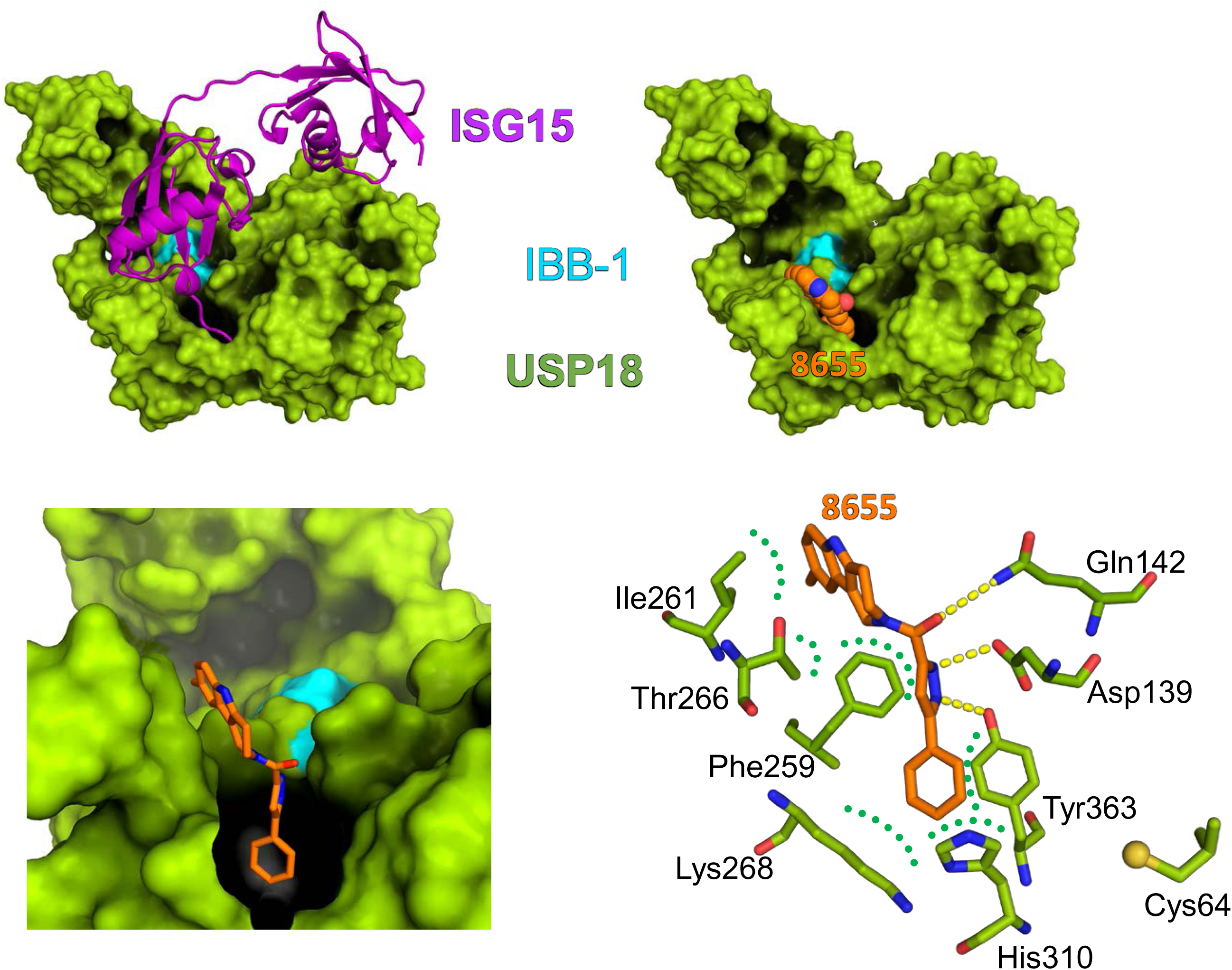
Structural model of the human USP18/ISG15 complex. USP18 is shown with solvent accessible surface in green. ISG15 is shown in magenta. The interaction plane IBB-1 is indicated in blue. The compound 8655 binds into the cleft that accommodates the C-terminus of ISG15 and close to the IBB-1 site. Compound 8655 interacts with hUSP18 via three hydrogen bonds (indicated as yellow dotted line) and hydrophobic interactions (indicated as dotted green lines).

## Discussion

PPIs are of central relevance in how molecular properties of polypeptides are translated into biological functions and how a cell can react towards specific stimuli. Concordantly, alterations in PPIs caused either by mutations, small molecules or posttranslational modifications are a central theme in multiple pathological processes. Although modulation of PPIs for therapeutic purposes has been proven to be challenging, it is an emerging topic with great potential to alter molecular functions previously thought to be “undruggable”. Encouragingly, several PPIs modulators are meanwhile either in clinical trials or even approved(36).

Suitable assay systems to efficiently, robustly and at least semi-quantitatively monitor PPIs are a prerequisite to understand structure-function relationships and consequences of (disease causing) protein variations. Here we developed assay systems for various components of the USP18 signaling hub, which is central to control type I IFN signaling and deISGylation. The cell-based NanoBRET assays have the advantage that PPIs can be monitored quantitatively in steady state or upon treatment (e.g by type I IFN stimulation). In the future, it will thus be possible to further elucidate on how IFN signaling affects PPIs within the IFNAR/USP18/STAT2/JAK1 complex. The assay can also readily be adapted to monitor consequences of mutations found in patients with regard to PPIs of USP18 or associated polypeptides and how such interactions are affected in various physiological or pathological conditions. Likewise, results from structural-functional analysis or newly identified interaction partners can easily be validated even if antibodies for immunoprecipitation approaches are not available.

Within this context it needs to be mentioned that the BRET assay has its intrinsic limitations that need to be carefully considered. First, the fusion of relatively bulky Tags (NLuc and HaloTag) harbors the risk of steric hinderance. Thus, several different fusion variants need to be evaluated and structural modelling helps to predict steric hinderance. Notably, a lack of NanoBRET signal is thus not necessarily an indication of a lack of interaction, but might origin from sterical hinderance caused by the tags. Furthermore, NanoBRET ratios from different interaction partners cannot be compared directly, as this readout is not an absolute measurement for interaction or binding capacity. Here biochemical methods, which allow to directly measure molecule interactions quantitatively, such as Surface Plasmon Resonance (SRP/Biacore) or isothermal titration calorimetry (ITC), are more appropriate.

Our results show that transferring the NanoBRET assay to a stable cell format allows to establish cell-based, screening compatible assays for PPI. To our knowledge this is the first documented proof of principle that such assays can successfully be applied to identify PPI-modulating small molecules in semi HTS. Cell-based systems will never reach the capacity of *in vitro* assays, which allow screening of thousands of small molecules. However, our results show that the combination of computational preselection tools with stable cell-based NanoBRET assay systems can successfully be applied to identify lead compounds interfering with PPIs. A clear advantage of cell-based screens is that only cell permeable compounds can affect the NanoBRET readout.

The work described here, led to the discovery of ZHAWOC8655, the first cell permeable small molecule that disturbs USP18/ISG15 binding and inhibits USP18 protease activity *in vitro*. Further medicinal chemistry optimization and pharmacological studies with this lead molecule are underway to develop this compound class towards chemical tools and ultimately clinical drug candidates.

The assay cell lines described here might also identify previously unknown endogenous proteins affecting the hSTAT2/hUSP18 or hISG15/hUSP18 interaction using siRNA or CRISPR screening libraries. Notably, it is also possible to monitor changes in the stability of one interaction partner. Within that context it is noteworthy that hUSP18 stability is affected by proteasomal degradation and that hISG15 is essential to secure its stability. Destabilizing hUSP18 by PROTACS for example could be a suitable strategy to enhance the IFN response and/or overcome desensitization, which might be useful for antiviral or IFN sensitive anti-tumor strategies. The cell lines developed here, efficiently allow to monitor protein levels of the NLuc-hUSP18 fusion protein and should thus also be applicable to identify or optimize hUSP18 targeting PROTACs. Likewise, identified compounds interfering with interactions could be tested for their molecular binding capacity to hUSP18 and allow selection of high affinity binding moieties for PROTAC construction.

The approach taken here is a suitable template applicable to monitor other PPIs and inhibitor screening approaches. The combination of small molecule preselection by *in silico* virtual screening and NanoBRET-based stable cell assays hold great potential for identifying PPI modulating compounds in various settings.

### Experimental procedures

#### Cloning of expression plasmids

For the transient NanoBRET assays, vectors were the HaloTag could be fused to the N-terminus (pFN21A) or the C-terminus (pFC14K) of a gene were used together with plasmids were the NLuc could be fused to the N-terminus (pFN31K) or the C-terminus (pFC32K) of the other gene of the PPI. Particularly, the coding sequence of hUSP18 and hSTAT2 were purchased from Promega and cloned into pFN31K and pFC14K vectors, respectively, with the Flexi® Cloning system (Promega) according to manufacturer’s instructions. To generate pFN31K hUSP18^ΔIBB1^ and hUSP18^C64A^ plasmids, coding sequences were amplified from pTriEx™2 hUSP18^ΔIBB1^ and pTriEx™2 hUSP18^C64A^ (13), respectively using the primer pair SgfI_hUSP18 (5’AAAAAAGCGATCGCCATGCCTGCTGTGGCTTCAG’3) and rev_hUSP18-PmeI (AAAAAAGTTTAAACTGTGGCTACATCAGTTACTCGTG). The coding sequence of PLpro was amplified from pET15b-PLpro vector (https://doi.org/10.1016/j.phymed.2023.155176) with primers SgfI_PLpro_fwd (5’TAAAGCGATCGCCATGGTGAGGACTATTAAGGTGTTTAĆ3) and PLpro_PmeI_rev (5’CTTAGTTTAAACTGCGGCCGCTATGGTTG’3) and inserted into vectors pFN21A, pFN31K, pFC14K and pFC32K with the same system. The plasmid pFN21A hISG15 was directly obtained from Promega.

To generate the stable cell lines, the same Flexi® Cloning system (Promega) was used to construct the NanoBRET BiBRET vectors. The coding sequences of hUSP18 and hUSP18^ΔIBB1^ were cloned into the pFN226C vector, resulting in N-terminal fusion with NLuc under the control of a minimal CMV promoter. The hISG15 sequence was inserted into the pFN222K vector, where it was fused to an N-terminal HaloTag and driven by a full-length CMV promoter. Similarly, hSTAT2 was cloned into the pFC221K vector, enabling C-terminal fusion with NLuc under the regulation of a full-length CMV promoter. To generate combined BiBRET vectors, containing both fusion-gene constructs, the vectors pFN226C hUSP18, pFN226C hUSP18^ΔIBB1^, pFN222K hISG15WT and pFC221K hSTAT2WT were digested with AscI and NotI-HF (NEB). Gel purification was performed to extract the fragments containing the CMV minimal promoter together with the coding sequence of hUSP18 or hUSP18^ΔIBB1^. These fragments were then cloned into pFN222K hISG15T and pFC221K hSTAT2 respectively using the Rapid DNA Dephos & Ligation kit (Roche) according to Manufacturer’s instructions. The correct cloning of all plasmids was confirmed by Sanger Sequencing carried out by Eurofins Genomics.

#### Generation of stable NanoBRET™ cell lines

HEK293 cells were transfected with AgeI-linearized combined BiBRET vectors pFN222K hISG15 hUSP18WT, pFN222K hISG15 hUSP18^ΔIBB1^ and pFC221K hSTAT2 hUSP18 using Fugene® HD transfection reagent (Promega) according to Manufacturer’s instructions. Stably transfected cells were selected by Blasticin S treatment (7 µg/mL) and single cell clones were picked by hand.

#### NanoBRET™ assay

Transient NanoBRET™ assay was performed according to Manufacturer’s instructions. Briefly, HEK293(T) cells were seeded at 4×10^5^ cells/mL and transfected with NanoLuc® and HaloTag® fusion vector at a ratio of 1:100 (2 µg of HaloTag and 20 ng of Nanoluc). Approximately 20 hours later, cells were harvested, resuspended in NanoBRET™ assay medium (phenol red-free Opti-MEM™, supplemented with 4% FCS) and seeded at 2×10^4^ cells (transient transfection) per well into white 96-well flat-bottom tissue culture-treated plates (Greiner) in the absence (0.1 % DMSO vehicle control) or presence of 100 nM HaloTag® NanoBRET™ 618 Ligand in triplicate. Immediately before measurement, the NanoBRET™ Nano-Glo® substrate (Promega) was added at 10 µM. Luminescence signals were measured at a wavelength range of 415 – 485 nm for the NLuc® luciferase donor and 610 – 700 nm for acceptor HaloTag® NanoBRET™ 618 Ligand with the Tecan Spark 10M plate reader.

The inhibitor GRL-0617 (Sigma-Aldrich) was dissolved in DMSO to reach a concentration of 10 mM. HEK293(T) cells were seeded at 4×10^5^ cells/mL and transfected with NanoLuc® and HaloTag® fusion vectors at a ratio of 1:100. Approximately 20 hours later, cells were harvested, resuspended in NanoBRET™ assay medium (phenol red-free Opti-MEM™, supplemented with 4% FCS) and seeded at 2×10^4^ cells per well into white 96-well flat-bottom tissue culture-treated plates (Greiner) in the absence (0.1 % DMSO vehicle control) or presence of 100 nM HaloTag® NanoBRET™ 618 Ligand in triplicate. Then different concentrations of GRL-0617 were prepared in phenol red-free Opti-MEM™ with the correspondent amount of DMSO and added to the cells for a final concentration range of 0,001-100 µM. 24h post-treatment, the NanoBRET™ Nano-Glo® substrate (Promega) was added at 10 µM. Luminescence signals were measured at a wavelength range of 415 – 485 nm for the NLuc® luciferase donor and 610 – 700 nm for acceptor HaloTag® NanoBRET™ 618 Ligand with the Tecan Spark 10M plate reader.

#### Optimization of the NanoBRET™ assay for screening applications

In preparation for using the NanoBRET™ assay for screening applications, the experimental assay setup was optimized. The assay was downsized from 96-well to a 384-well format. The liquid handling procedures and machine setup were optimized to ensure efficient and economical workflow. In addition, the test conditions for compound screening were optimized to determine the best pipetting procedures for compound dilution as well as cell treatment conditions. NanoBRET™ test cells (HEK293 cells transiently or stably transfected with the NanoBRET™ PPI pairs) and test compounds were prepared and mixed at a ratio of 1:1 in the absence (vehicle control – 0.1 % DMSO) or presence of 100 nM HaloTag® ligand 618 (Promega). The final concentration of DMSO within the assay medium amounted to 2.1 %. Final cell concentration was 2×10^5^/mL. For high-throughput measurements the cell suspension was plated into white 384-well tissue culture-treated flat-bottom plate at 40 µL per well in triplicate using the Freedom Evo 200 Liquid Handling System with disposable tips (Tecan) and incubated for approximately 24 hours at 37°C prior to measurement. The NanoBRET™ Nano-Glo® substrate (Promega) was added to the plate with the Multiflo Microplate Dispenser (Biotek). The plate was centrifuged for approximately 20 s at 300 xg. Luminescence measurements were performed with the Synergy H4 Hybrid reader (Biotek) using the following settings: emission 460/40 nm and a gain of 135 for the donor signal, emission 620/40 and a gain of 180 for the acceptor signal.

#### Data analysis

To calculate the corrected NanoBRET™ ratio (mBU) the acceptor signal was divided by the donor signal and the background signal (DMSO vehicle) was subtracted from the experimental sample (HaloTag® NanoBRET™ 618 Ligand). NanoBRET™ ratios of test compound treated samples were normalized to the vehicle control sample. Data depiction and statistical analysis were performed using GraphPad Prism 9.3. A P value of less than 0.05 was considered statistically significant (*P < 0.05; **P < 0.01; ***P < 0.001; ****P < 0.0001). The dose-response curve of each compound was graphed with Graphpad Prism 9.3 software using the curve fit function with three parameters setting utilizing the following equation: Y=Bottom + (Top-Bottom)/(1+(X/IC50)).

#### Virtual screening and docking calculations

For virtual screening, compound libraries from Enamine and Life Chemicals comprising in total 60,000 molecules were used. The structure of murine USP18 (PDB code: 5chv) served as receptor in high-throughput docking calculations using AutoDock Vina (31). In order to test *in silico* binding of selected compounds to human USP18, we generated models using RoseTTAFold (https://www.science.org/doi/10.1126/science.abj8754) and Alphafold (https://doi.org/10.1038/s41586-021-03819-2) and checked those for agreement. A consistent model was selected for docking calculations, which were performed with the programs AutoDock Vina and PLANTS (https://doi.org/10.1021/ci800298z). The results with highest docking scores and similar poses were selected. 3D models of the NLuc and HaloTag fusion proteins were generated with AlphaFold and arranged for the figure using Coot (https://doi.org/10.1107/s0907444910007493). The figures were generated using PyMol.

#### Determination of inhibition constants

A synthetic cDNA encoding human USP18 residues 52-372 with an N-terminal hexahistidine-tag and synthetic cDNA encoding human ISG15-C78S were cloned into pET-Duet vector via NcoI and BamHI for USP18 and NotI and XhoI for ISG15, respectively. Human USP18 was heterologously expressed in *Escherichia coli* BL21(DE3) and purified via Ni^2+^-immobilized affinity chromatography using a HisTrap FF Crude column (Cytiva) as described by the manufacturer.

Enzyme activity was measured using 8 nM USP18 in 20 mM HEPES pH 7.4, 10 mM NaCl, 150 mM KCl, 5 mM DTT, 0.2 mg/ml BSA, 5% glycerol with 100 nM C-terminal domain of human ISG15 conjugated to the fluorophore rhodamine (ISG15ct-RhoMP) as a substrate. The increase in fluorescence at 535 nm was followed using an excitation wavelength of 458 nm. All measurements were performed at 293 K in triplicates.

Inhibitor ZHAWOC8655 was added to the assay at concentrations 0–30 µM and incubated for 5 min before starting the assay by addition of ISG15ct-RhoMP at a final concentration of 100nM. The dose response curve for IC_50_ determination was calculated with OriginPro, Version 2020. (OriginLab Corporation, Northampton, MA, USA).

#### Analysis of protein expression

HEK293 cells were transiently transfected with plasmids encoding for NLuc-hUSP18, HaloTag-hISG15 and hSTAT2-HaloTag using Fugene® HD transfection reagent (Promega) according to manufacturer’s instructions. Cells were lysed after 48 hours with a lysis buffer containing 50 mM Tris-HCl pH7.4, 150 mM NaCl, 1 mM EDTA, 1 % (v/v) Triton X-100. Alternatively, cells stably expressing the NanoBRET interaction pairs NLuc-hUSP18/HaloTag-hISG15 and NLuc-hUSP18/hSTAT2-HaloTag were lysed with the same lysis buffer. The proteins were denatured in Laemmli buffer containing 0.1 M DTT and incubated at 95°C for 5 min prior to Immunoblot analysis with antibodies directed against NLuc (Promega), HaloTag (G928A, Promega), hUSP18 (4813, Cell Signaling) and GAPDH (AB2302, Millipore).

## Acknowledgement

This work was supported by the Deutsche Forschungsgemeinschaft (DFG) grants 423813989/GRK2606 and SFB/TRR 167 Project number 259373024 to KPK, by ZHAW Anschubfinanzierung (start-up funding) to RR and by The Swiss National Science Foundation (SNSF) grant CRSK-3_221176 / 1 to RR. Further support to KPK was provided from the DFG under Germany’s Excellence Strategy (CIBSS–EXC-2189–Project ID 390939984).

