## Supplemental Figures 1 to 3 for "A NanoBRET-based assay monitoring interactions in the USP18 signaling hub identifies the first cell-penetrant small molecule compromising USP18/ISG15 binding"

### Suppl Figure 1: BiBRET Vector Card pFN226C

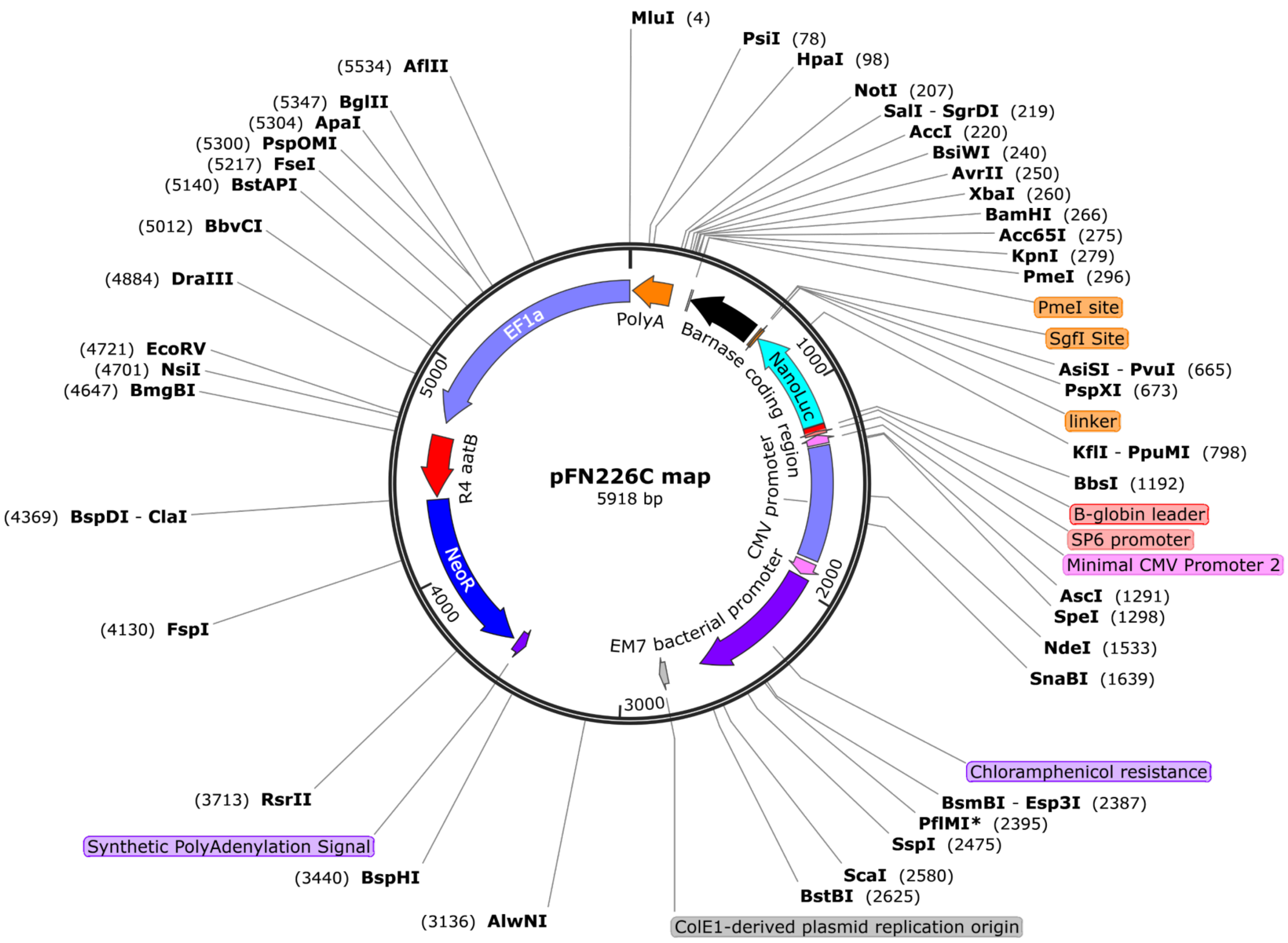

**Suppl Figure 2: BiBRET Vector Card pFN222K**

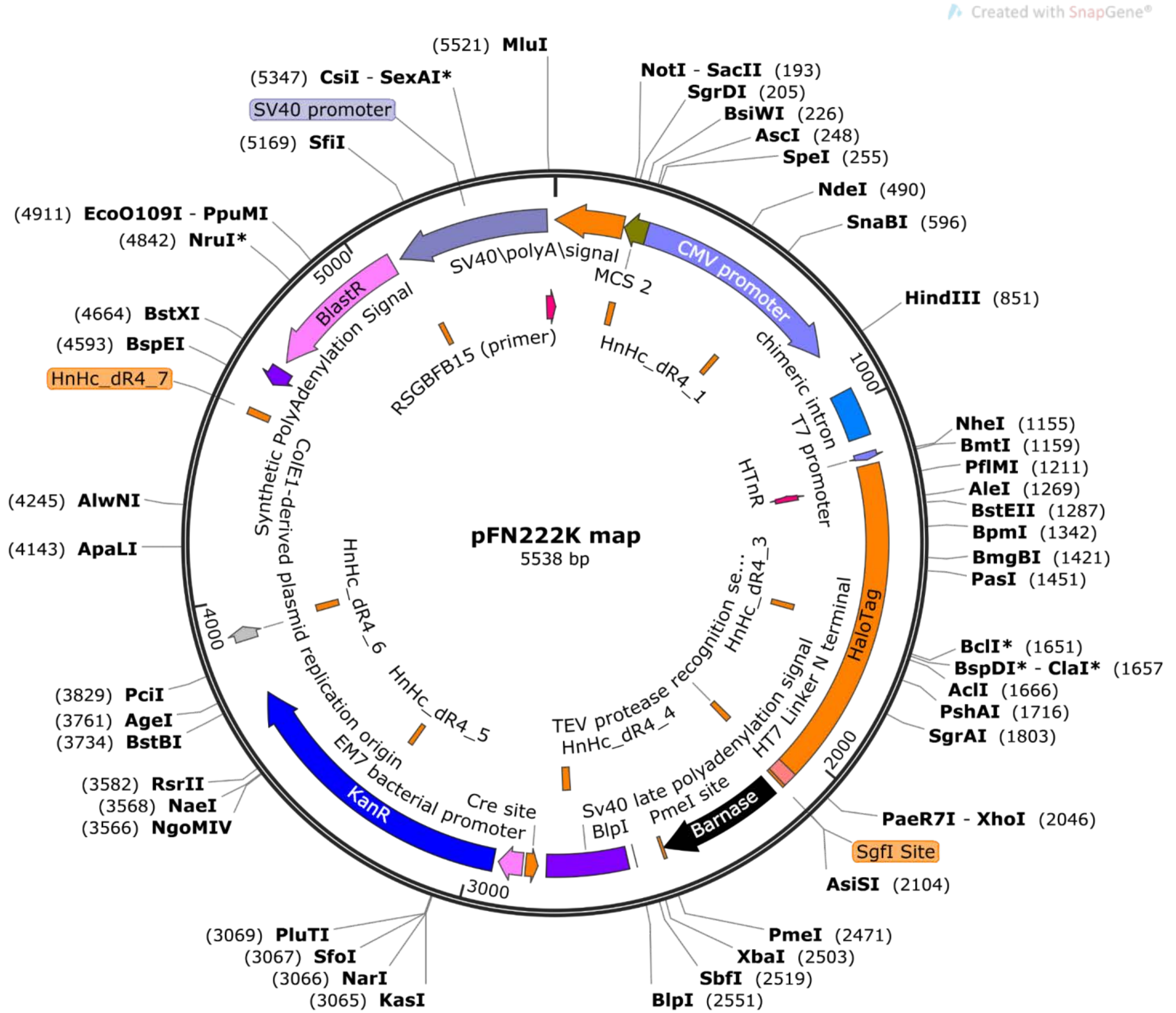

### Suppl Figure 3: BiBRET Vector card pFC221K

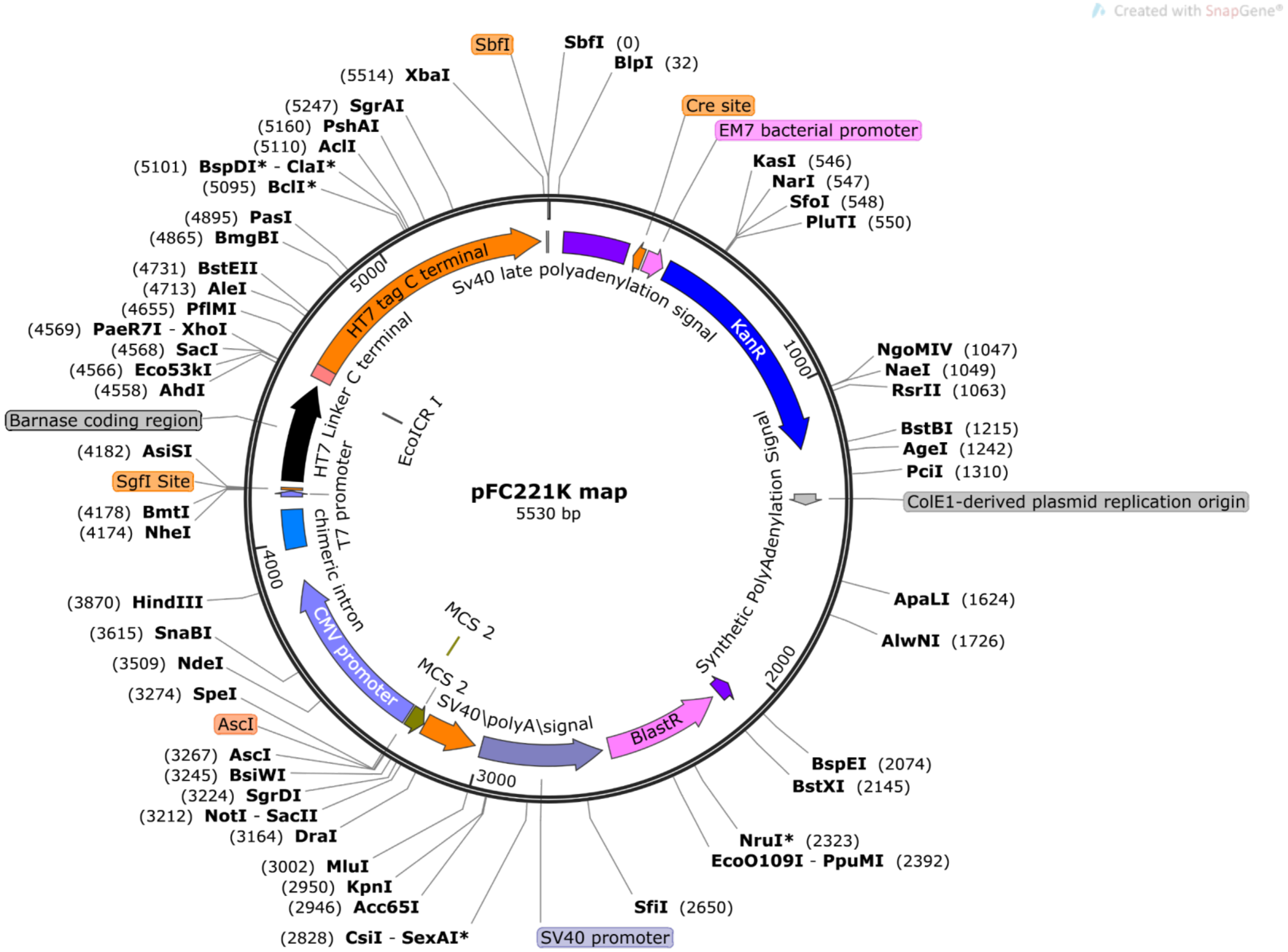
